# Global detection of DNA repair outcomes induced by CRISPR-Cas9

**DOI:** 10.1101/2021.02.15.431335

**Authors:** Mengzhu Liu, Weiwei Zhang, Changchang Xin, Jianhang Yin, Yafang Shang, Chen Ai, Jiaxin Li, Fei-long Meng, Jiazhi Hu

**Author notes:** These authors contributed equally to this work (Mengzhu Liu, Weiwei Zhang, Changchang Xin).

## Abstract

CRISPR-Cas9 generates double-stranded DNA breaks (DSBs) to activate cellular DNA repair pathways for genome editing. The repair of DSBs leads to small insertions or deletions (indels) and other complex byproducts, including large deletions and chromosomal translocations. Indels are well understood to disrupt target genes, while the other deleterious byproducts remain elusive. We developed a new *in silico* analysis pipeline for the previously described primer-extension-mediated sequencing assay to comprehensively characterize CRISPR-Cas9-induced DSB repair outcomes in human or mouse cells. We identified tremendous deleterious DSB repair byproducts of CRISPR-Cas9 editing, including large deletions, plasmid integrations, and chromosomal translocations. We further elucidated the important roles of microhomology, chromosomal interaction, recurrent DSBs, and DSB repair pathways in the generation of these byproducts. Our findings provide an extra dimension for genome editing safety besides off-targets. And caution should be exercised to avoid not only off-target damages but also deleterious DSB repair byproducts during genome editing.

## Introduction

Genome editing technologies based on engineered nucleases not only greatly change the way we study life sciences but also cast light on the treatment of human genetic diseases (Doudna 2020; Wang et al. 2020). Among these powerful editing toolboxes, the clustered regularly interspaced short palindromic repeat (CRISPR) and associated protein (Cas) engineered from the bacterial defense system are the most widely used ones. The *Streptococcus pyogenes* Cas9 (referred to as Cas9 hereafter) from type II CRISPR-Cas systems is the earliest Cas protein to be engineered for performing genome editing in human cells (Cong et al. 2013; Jinek et al. 2013; Mali et al. 2013). CRISPR-Cas9 is a two-component editing system, comprising of a Cas9 protein with cleavage activity and a guide RNA (gRNA) to bind both Cas9 and target DNA (Jinek et al. 2012). CRISPR-Cas9 is in principle able to induce doublestranded DNA breaks (DSBs) at any locus 3 base pairs (bp) upstream of an NGG protospacer adjacent motif (PAM). Besides inducing mutations at target sites, CRISPR-Cas9 may also generate unintended damages at homologous off-target sites, raising great safety concerns. These off-target activities of CRISPR-Cas9 can be largely minimized by using high-fidelity Cas9 variants, choosing a better target sequence instead, or rapid activity shut-off by anti-CRISPR (Pawluk et al. 2018; Anzalone et al. 2020; Hendriks et al. 2020).

The first step for CRISPR-Cas9 editing is to initiate DSBs at DNA target sites that are complementary to the gRNAs. The endogenous DNA repair pathways are subsequently activated to create a variety of DNA repair outcomes, including a large number of insertions and deletions. There are two main DSB repair pathways in mammalian cells, non-homologous end joining (NHEJ) and homologous recombination (HR). To repair Cas9-induced DSBs, NHEJ directly fuses two broken ends to seal DSBs, frequently accompanied by small insertions or deletions (indels) that are less than 20 bp; while HR requires external homologous donor DNA to introduce intended mutations (Yeh et al. 2019). Besides, the alternative end-joining (A-EJ), also termed as microhomology-mediated end joining (MMEJ), is also involved in DSB repair after the exposure of microhomologies at juxtaposed broken ends following end processing (Alt et al. 2013; Sfeir and Symington 2015). MMEJ requires microhomologies that range from 2-20 bp while NHEJ might also utilize microhomologies less than 4 bp (Truong et al. 2013; Chang et al. 2017). Both NHEJ and MMEJ are error-prone and thereby may generate deleterious DSB repair byproducts, including large chromatin deletions and chromosomal translocations, resulting in chromosomal abnormality or tumorigenesis (Alt et al. 2013; Chang et al. 2017; Zhao et al. 2020). In this context, large deletions, chromosomal translocations, or even chromosome loss has been detected by different research groups in mouse and human stem cells after CRISPR-Cas9 editing (Zuo et al. 2017; Adikusuma et al. 2018; Kosicki et al. 2018; Cullot et al. 2019; Zuccaro et al. 2020).

Approaches to manipulate DSB repair pathways have been developed to enhance genome editing (Yeh et al. 2019; Ling et al. 2020). For instance, inhibitors for key NHEJ factors KU or LIG4 are used to increase the incorporated rate of donor fragments by enhancing HR (Chu et al. 2015; Maruyama et al. 2015; Hu et al. 2018; Riesenberg and Maricic 2018); RAD51 has been fused with Cas9 nickase to facilitate the insertion of doublestranded oligonucleotide (Rees et al. 2019). In contrast, inhibitors for CtIP or RAD52 suppress HR and promote single-stranded oligonucleotide-mediated editing (Riesenberg and Maricic 2018). However, blocking the DSB repair pathway would alter the spectrum of CRISPR-Cas9-induced repair outcomes and may threaten genome integrity. For this reason, the danger of using inhibitors for key DNA repair factors on genome integrity during genome editing remains to be elucidated.

Comprehensive assessment of global DSB repair outcomes would facilitate our understanding of the origins of the deleterious byproducts including large deletions and chromosomal translocations as well as help improve genome-editing safety. Here, we developed a new *in silico* analysis pipeline to identify genome-wide DSB repair outcomes based on the high-throughput sequencing data generated via previously described primerextension-mediated sequencing (PEM-seq) (Yin et al. 2019). We find that large deletions heavily depend on microhomologies and large insertions contain substantial vector integrations. Chromosomal translocations distribute widely in the genome and are often dominated by off-target or other recurrent DSBs. Furthermore, we also detect an increased level of chromosomal abnormality in the absence of the NHEJ repair pathway.

## Results

### Detecting global DNA repair outcomes of CRISPR-Cas9 by PEM-Q pipeline

To gain insight into the full spectrum of DNA repair products resulted from genome editing exerted by CRISPR-Cas9, we have developed the PEM-seq to capture unknown broken end (prey) fused with the target DSBs (bait) in cells 72 hours post-transfection (Fig. 1A). The identified repair outcomes are further categorized into re-joinings of the target broken ends, leading to insertions and deletions, and intra- or inter-chromosomal translocations (Fig. 1A). In order to quantify indels that are invisible to the previous SuperQ pipeline (Yin et al. 2019), we employed *bwa-mem* instead of *bowtie2* for genome alignment and optimized the *in silico* analysis flow to develop a new pipeline, termed as PEM-Q (Supplemental Fig. S1A; see Methods for details). Then we used PEM-Q to analyze the deep sequencing data from CRISPR-Cas9-edited K562 cells at *HBB* locus in parallel with CRISPResso (Pinello et al. 2016). The distribution pattern of indels identified by PEM-Q was almost identical to that identified by CRISPResso (Fig. 1B). The SuperQ and high-throughput genome-wide translocation sequencing (HTGTS) pipelines have been used to identify translocation junctions and off-targets of CRISPR-Cas9 in HEK293T cells at various loci (Frock et al. 2015; Hu et al. 2016; Yin et al. 2019). We re-analyzed the same sequencing data from *RAG1* locus by PEM-Q and found highly similar genome-wide translocation patterns among these pipelines (Fig. 1C). Moreover, PEM-Q identified 7 more off-targets than SuperQ and 11 more off-targets than HTGTS, showing a higher sensitivity of detecting off-targets (Supplemental Fig. S1B and S1C). Therefore, PEM-Q is a unified *in silico* analysis pipeline for detecting global DNA repair outcomes of CRISPR-Cas9.

**Figure 1.**
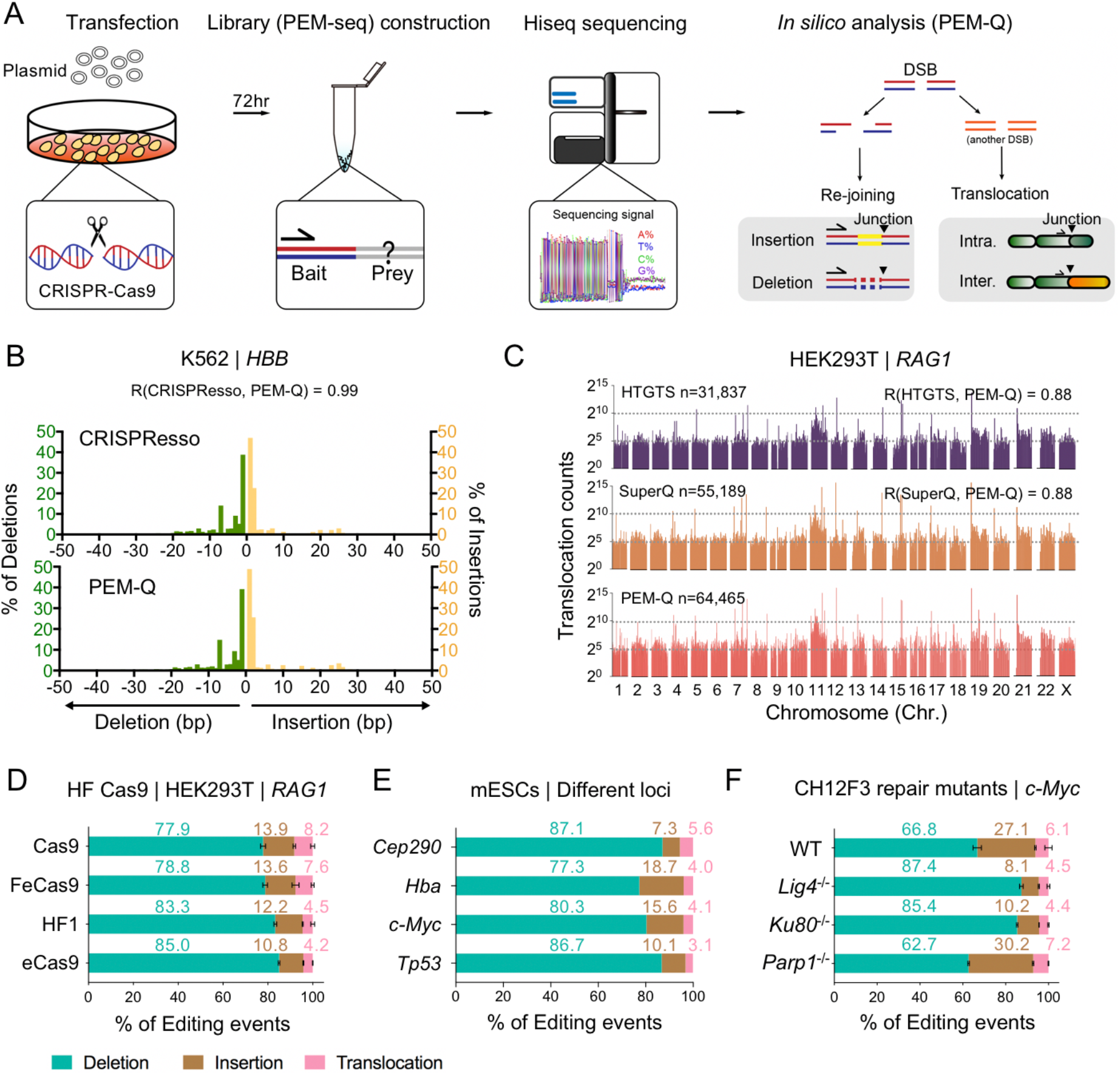
Detecting global DNA repair outcomes of CRISPR-Cas9 by PEM-Q pipeline. **(A)** Experimental procedures. Sequencing libraries were generated by PEM-seq with Hi-seq sequencing. The identified re-joining events contain insertions and deletions, while translocation junctions beyond upstream and downstream 500 kb from the target site in the target chromosome are defined as intra-chromosomal translocations (Intra.) and junctions from different chromosomes are inter-chromosomal translocations (Inter.) Black triangles represent identified junctions.**(B)** Reanalysis the published data (Pinello et al., 2016) by PEM-Q to compare with CRISPResso at the *HBB* locus in K.562 cells. Green bars represent the frequency of deletions at indicated length normalized to total deletions; while yellow bars represent insertions. Pearson correlation (R) between CRISPResso and PEM-Q is 0.99. **(C)** Re-analysis of PEM-seq data (Yin et al., 2019) by PEM-Q versus HTGTS and SuperQ at the *RAG1* locus in HEK293T cells. Total translocation junctions from three replicates are shown with 2-Mb bins on a log scale. Total numbers (n) of translocations and Pearson correlation (R) are indicated. **(D-F)** Bar charts showing percentages of deletions (cyan), insertions (brown) and translocations (pink). (D) High-fidelity (HF) Cas9 enzymes at *RAG1* locus in HEK293T cells. **(E)** Different target loci *(Cep290, Hba, c-Myc* and *Tp53}* in mESCs. **(F)** The *c-Myc* locus in CH12F3 cells with indicated backgrounds. Error bars, mean ± SD for **D** and **F**.

We next employed PEM-Q to systematically analyze public PEM-seq data from CRISPR-Cas9-edited HEK293T cells at the *RAG1* gene (Yin et al. 2019).We divided the repair outcomes into re-joinings and translocations for further analysis. The re-joinings of the target DSB result in deletions and insertions, while chromosomal translocations are derived from the fusion of target DSB with another DSB either in the same chromosome or other chromosomes (Fig. 1A). The PEM-Q-identified deletions, insertions, or translocations were highly reproducible from three repeat libraries (Supplemental Fig. S1D-F). Furthermore, deletions were the main repair outcomes concentrated in the upstream and downstream 15 base pairs (bp) around the cleavage site, approximately 78% of total editing events; insertions were about 14%, enriched within a 5-bp region around the cleavage site; while translocations distributed widely in the genome at a rate of 7-9% (Supplemental Fig. S1D-F). A number of high-fidelity variants have been developed to reduce the off-target activity of Cas9 (Yin et al. 2019; Hendriks et al. 2020). We re-analyzed the PEM-seq data of three high-fidelity variants eCas9, HF1, and FeCas9 versus Cas9 from HEK293T cells at the *RAG1* gene (Yin et al. 2019). High-fidelity variants showed similar levels of different repair outcomes, including high levels of translocations (Fig. 1F), suggesting the inadequate ability of high-fidelity Cas9 variants to reduce deleterious DNA repair byproducts.

We also used CRISPR-Cas9 to target the *Cep290, Hba, c-Myc,* and *Tp53* loci in mouse embryonic stem cells (mESCs) and then generated PEM-seq libraries for PEM-Q analysis. Repair outcomes in mESCs showed similar compositions as those in HEK293T cells despite the percentages of different repair outcomes varied at examined target sites (Fig. 1E). Besides, we performed CRISPR-Cas9 editing at the *c-Myc* and *Bcr* locus in the mouse CH12F3 B cells with different repair backgrounds. The deficiency of core NHEJ factors Ku80 or DNA Ligase 4 (Lig4) induced a significant increase of deletions, while Parp1-deficient cells displayed a similar pattern to that of wild-type (WT) cells (Fig. 1F and Fig. S1G). Collectively, these data suggest that Cas9 repair-outcomes are nonrandom as to be predictable, so we decided to further explore these repair outcomes of CRISPR-Cas9 with comprehensive analysis capability of PEM-Q.

### Prevalent microhomologies at large deletions of CRISPR-Cas9

Deletional rejoining identified by PEM-Q was the most abundant repair outcome after CRISPR-Cas9 editing even when normalized to the total sequencing events (Fig. 2A and Supplemental Fig. S2A). Chromosomal deletions were widely distributed downstream of the cloning primer binding site and expanded as long as hundreds of kb at the *c-Myc* locus while tens of kb at the *Bcr* locus in CH12F3 cells, depending on the cutting efficiency at two loci (Fig. 2B and Supplemental Fig. 2B), consistent with previous reports (Frock et al. 2015; Yin et al. 2019). In this context, we divided Cas9-induced deletions into two parts: small deletions within 100 bp and large deletions larger than 100 bp. Small deletions were the main deletional events at a percentage of more than 94%, while large deletions were also unignorable, more than 5% at both *c-Myc* and *Bcr* loci (Fig. 2B, 2C, and Supplemental Fig. S2B). Specifically, 0.8% and 0.3% deletions were larger than 3 kb at the *c-Myc* or *Bcr* locus, respectively (Fig. 2B and Supplemental Fig. S2B). Similar findings were obtained in four different target sites in mESCs, and, notably, large deletions were increased to 11.7% and 14.7% at the *c-Myc* and *Hba* loci, respectively (Supplemental Fig. S2C).

**Figure 2.**
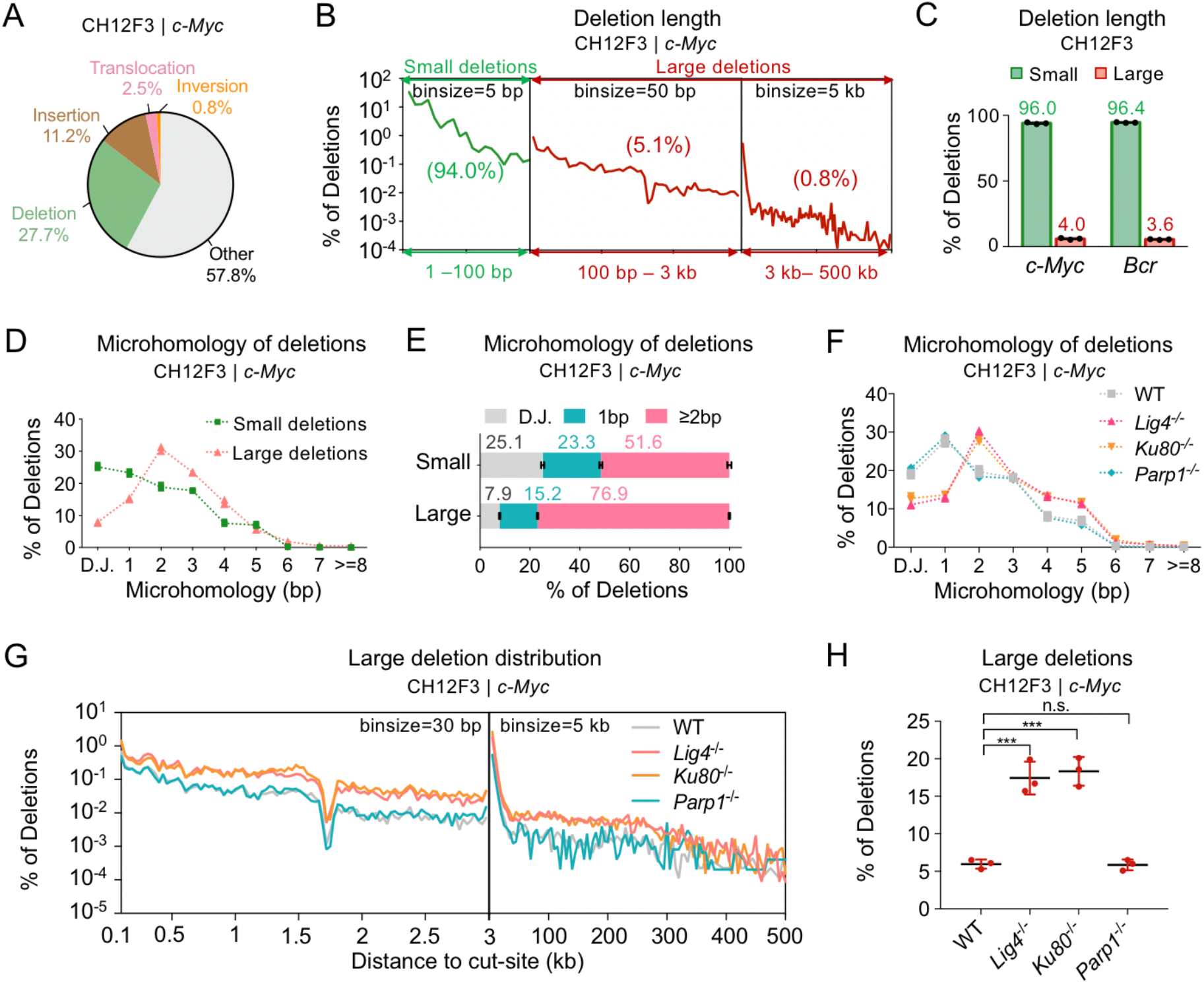
Prevalent microhomologies at large deletions of CRISPR-Cas9. **(A)** Pie chart of total sequencing reads at the *c-Myc* locus in CH12F3 cells. Total sequencing reads include deletions, insertions, translocations, inversions, and other reads (including germline sequences). The average percentages of three repeats are shown. (B) The distribution pattern of deletions at the *c-Myc* locus in CH12F3 cells. Total junctions of three repeats are plotted on a log scale. Percentages of deletions within each region are shown in the brackets. Please note that 5 bp, 50 bp, and 5 kb bin-sizes are used for the three regions, respectively. (C) Percentages of small and large deletions at the *c-Myc* and *Bcr* loci in CH12F3 cells. Error bars, mean ± SD. **(D** and **E)** Line plot **(D)** and bar chart **(E)** of microhomologies with indicated length in small or large deletions at the *c-Myc* locus in CH12F3 cells. Only deletions cross the cut-sites are used for analysis. D.J., direct joining. Error bars, mean ± SD. **(F)** Line plot of microhomologies with indicated length in total deletions at the *c-Myc* locus in CH12F3 cells with indicated backgrounds. (G) The distribution patterns of large deletions at the *c-Myc* locus in CH12F3 cells with indicated backgrounds. Please note that 30 bp and 5 kb bin-sizes are used for two regions, respectively. (H) Percentages of large deletions at the *c-Myc* locus in CH12F3 cells with indicated backgrounds. One-tailed t-test, ***, p < 0.0005; n.s., not significant. Error bars, mean ± SD.

Given that the formation of deletion requires end processing that promotes microhomologies for MMEJ, we next examined the usage of microhomology in these deletional events. Approximately 25% of small deletions preferred direct joining and only around half of them used microhomologies longer than 2 bp with a decreasing trend over length (Fig. 2D and 2E). Different from small deletions, large deletions heavily depended on microhomology and over 76% of large deletions used microhomology longer than 2 bp while direct joinings were only about 7.9% (Fig. 2D and 2E), consistent with previous findings (Owens et al. 2019). Similar findings were obtained in CRISPR-Cas9-edited CH12F3 cells at the *Bcr* locus and mESCs at four different loci (Supplemental Fig. S2D and S2E). For further verification, we examined the microhomology usage in CH12F3 cells deficient for Ku80 or Lig4. NHEJ predisposes to direct joining and is suppressed in Ku80- or Lig4-deficient cells (Chang et al. 2017). As anticipated, we detected an elevated level of microhomology usage in Cas9-induced deletions from Ku80-or Lig4-deficient CH12F3 cells (Fig. 2F and Supplemental Fig. S2F). Correspondingly, the level of large deletions from 100 bp to 300 kb was increased significantly in Ku80- or Lig4-deficient background (Fig. 2G and 2H).

However, the deficiency for Parp1 resulted in no significant effect on the deletion pattern at *c-Myc* locus despite a minor declined level of large deletions at the *Bcr* locus in CH12F3 cells (Fig. 2G, 2H and Supplemental Fig. S2G, S2H).

### Predictable small insertions and deleterious plasmid integrations

We also divided CRISPR-Cas9-induced insertions identified by PEM-Q into two parts: small insertions less than 20 bp and large insertions for 20 bp or more. Of note, repair outcomes with simultaneous insertions and deletions are categorized into insertions but not deletions in this study. Small insertions were 96.6% of all identified Cas9-induced insertions and 1-bp insertion dominated all insertions in CH12F3 cells at the *c-Myc* locus (Fig. 3A and 3B). Dominant small insertions were also detected in CRISPR-Cas9-edited CH12F3 cells at *Bcr* locus and mESCs at four different loci (Fig. 3B). In small insertions, the 1-bp insertions identical to the 4^th^ nucleotide T upstream of NGG occurred most frequently, up to 81% of total insertions at the *c-Myc* locus (Fig. 3C, #1 in top panel), resulted from the staggered cleavage of Cas9 (Shou et al. 2018; Chakrabarti et al. 2019). In this context, we found more examples of inserted nucleotides between the 4^th^ and 5^th^ nucleotides upstream of NGG, ranking high in all small insertions (Fig. 3C, #3, #6, and #10 in top panel). In the absence of Ku80 and Lig4, the 1-bp insertions declined significantly, especially for the top T insertions, from 81% in WT cells to 4.5% and 13.1% in Lig4- and Ku80-deficient cells, respectively (Fig. 3C-E and Supplemental Fig. S3A). Conversely, the level of insertions longer than 1 bp increased dramatically in both Lig4- and Ku80-deficient cells (Fig. 3D and 3E). Whereas in the absence of Parp1, both frequency and order of top 10 insertions were highly similar to those in WT cells. The reproducible patterns suggest that the sequence and frequency of small insertions are predictable with an unexplored mechanism.

**Figure 3.**
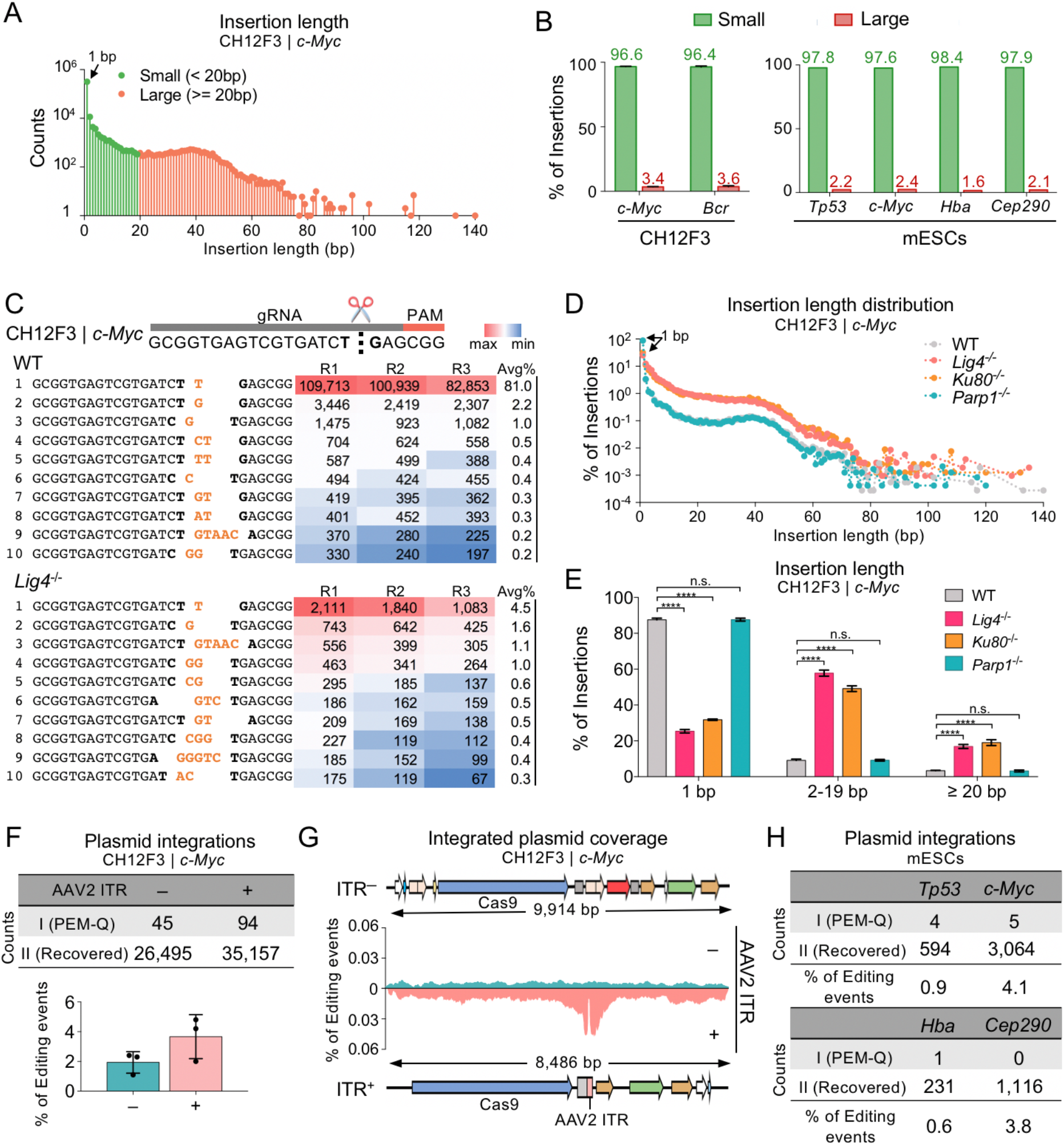
Predictable small insertions and deleterious plasmid integrations. **(A)** The distribution of insertions with indicated length at the *c-Myc* locus in CH12F3 cells. Total junctions of three repeats are plotted on a log scale. 1-bp insertion is indicated by the black arrow. (B) Percentages of small (< 20 bp) and large (≥ 20 bp) insertions at the *c-Myc* and *Bcr* locus in CH12F3 cells (left) and four indicated loci in mESCs (right). Error bars, mean ± SD. **(C)** Top 10 most frequent insertions at the *c-Myc* locus in WT and *Lig4*^-/-^ CH12F3 cells. The target site is shown on the top. Bases at cut-site are in bold and inserted bases are in orange. The numbers from three repeats (Rl, R2, R3) and average percentages (Avg%) of each type of insertions are listed in the table, filled with gradient color from the maximum (red) to the minimum (blue) frequencies. (D) The distribution patterns of insertion length at the *c-Myc* locus in CH12F3 cells with indicated backgrounds. Total junctions of three repeats are plotted on a log scale. 1-bp insertions are pointed out by the black triangles. (E) Bar chart showing the percentages of insertions with indicated length in CH12F3 cells with indicated backgrounds. Error bars, mean ± SD. One-tailed t-test, ****p < 0.0001; n.s., not significant. (F) The numbers of plasmid integrations with (+) or without (-) AAV2 ITR at the *c-Myc* locus in CH12F3 cells (top). Frequencies of integrated plasmid fragments are shown at the bottom. Error bars, mean ± SD. (G) Coverage of plasmid integrations at the *c-Myc* locus in CH12F3 cells with (bottom) or without (top) AAV2 ITR. **(H)** The numbers of plasmid integrations at indicated loci in mESCs.

We also noticed a pileup of large insertions around 40 bp in CRISPR-Cas9-edited CH12F3 cells at the *c-Myc* locus (Fig. 3A). We extracted the inserted sequences from CH12F3 cells at the *c-Myc* locus to align to the Cas9-carrying plasmid and found 45 distinct inserted sequences originated from the transfected plasmids (Fig. 3F, Structure I; Table S1), indicating potential plasmid integrations into the genome during CRISPR-Cas9 editing. To gain deep insight into plasmid integrations, we performed PEM-Q analysis against the mouse genome and then plasmid backbone sequence for sequence alignment. Three types of plasmid integrations with no overlap between the R1 and R2 sequencing reads were recovered at a frequency of ~2% of total editing events at the *c-Myc* locus (Fig. 3F and Supplemental Fig. S3B, Structure II). The inserted sequences evenly covered the whole plasmid backbone (Fig. 3G, top in cyan).

Interestingly, when using another Cas9-carrying plasmid with an adeno-associated virus 2 (AAV2) inverted terminal repeat (ITR) sequence for transfection, we observed an elevated level, although not statistically significant, of total plasmid integrations (Fig. 3F and Supplemental Fig. S3C). The ITR region forms a hairpin structure that affects the vector stability (Hanlon et al., 2019) and, therefore, the ITR regions become a hotpot for plasmid integration (Fig. 3G, bottom in salmon). Moreover, the ITR-carrying plasmid is integrated into the genome at the CRISPR-Cas9-edited mESCs at different target sites (Fig. 3H). This finding had striking similarities with viral integrations when using AAV to deliver Cas9 (Hanlon et al. 2019; Nguyen et al. 2021), which indicates that ITR or similar fragile sites is a significant cause for vector integrations.

### Distribution profile of translocations induced by CRISPR-Cas9

Translocation links the bait DSBs at the target site to genome-wide prey DSBs and thereby the translocation junctions represent the breakpoints of prey DSB in PEM-seq (Yin et al. 2019). Since large deletions can expand to the downstream region as long as 500 kb as revealed above (Fig. 2B), we excluded identified junctions within upstream and downstream 500 kb of target sites for translocation analysis. Translocation junctions distributed widely in the genome with an obvious enrichment at the target chromosome Chr15 when editing the CH12F3 cells at *c-Myc* locus via CRISPR-Cas9 (Fig. 4A). Similar enrichment in the target chromosome was also detected in PEM-seq libraries from CRISPR/Cas9-edited mESCs or human HEK293T cells at various loci (Fig. 4B and 4C). Translocation requires the proximity of two DSBs and chromatin interaction plays an indispensable role in the formation of translocation as revealed by the I-SceI-induced translocations (Zhang et al. 2012). We employed the 3C-HTGTS (Jain et al. 2018) to check the global interactions with the Cas9-target site. The distribution profile of translocation junctions was highly correlated with the interaction intensity revealed by 3C-HTGTS globally or within the target chromosome (Fig. 4D and Supplemental Fig. S4A). With this regard, the target chromosome Chr15 showed the most robust interaction intensity with the target site and thereby had the most translocations (Fig. 4D). We also used 5-Gy ionizing radiation (IR) to generate genome-wide DSBs that is independent of Cas9. IR-induced translocations captured by Cas9-induced target DSBs at the *c-Myc* locus were also correlated to interaction intensity (Fig. 4D).

**Figure 4.**
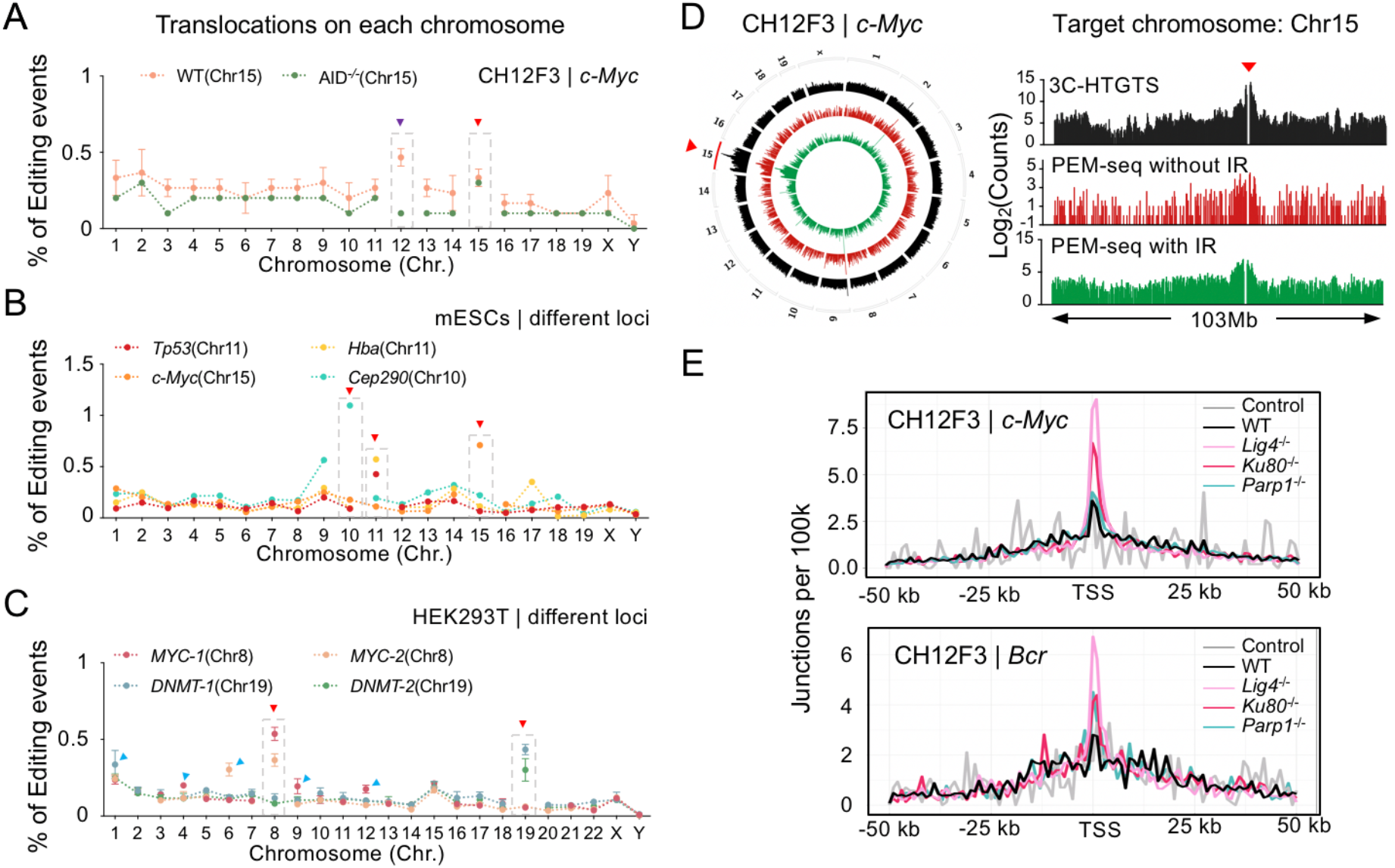
Translocations induced by CRISPR-Cas9 editing. **(A)** The distribution patterns of translocation junctions on each chromosome at the *c-Myc* locus in WT and AID^-/-^ CH12F3 cells. The target chromosome Chrl5 is highlighted by dashed-line boxes and indicated by the red triangle. Chrl2 is indicated by a purple triangle. (B) The distribution patterns of translocation junctions at indicated loci in mESCs. **(C)** The distribution patterns of translocation junctions at indicated loci in HEK293T cells. Chromosomes harboring robust off-target sites are pointed by blue triangles. (D) The distribution patterns of identified junctions in the entire genome (circos plot) or Chrl5 (bar graph) by 3C-HTGTS (black), PEM-seq with or without (red) IR at the *c-Myc* locus in AID ‘^-^CH12F3 cells. Signals were binned into 2 Mb intervals and plotted on a log scale. The upstream and downstream 500 kb region of the *c-Myc* locus (indicated by a red triangle) is removed. (E) The distribution patterns of translocation junctions around TSSs in CH12F3 cells with indicated backgrounds at the *c-Myc* locus (top) or *Bcr* locus (bottom). Translocations within the *IgH* region are excluded for analysis. Control represents primer control libraries without Cas9 cutting.

Chr12 also exhibited an enrichment of translocations in the CRISPR-Cas9-edited CH12F3 cells at the *c-Myc* locus (Fig. 4A). CH12F3 cells can undergo class switch recombination stimulated by anti-CD40/Interleukin 4/ TGF-β and activation-induced deaminase (AID) initiates substantial DSBs in the switch (S) regions (Liu et al. 2020). We examined the hotspot region in Chr12 and all the DSBs enriched at S regions as anticipated (Supplemental Fig. S4B). Moreover, the knock-out of AID resulted in a fall back of translocation level in Chr12 (Fig. 4A, in green). S regions are only activated in B lymphocytes, however, recurrent DSBs often occur at off-target sites during genome editing in non-lymphocytes. We checked the translocation junctions in CRISPR-Cas9-edited HEK293T cells and detected an elevated level of translocations in chromosomes harboring robust off-target sites (Fig. 4C). Besides off-target sites, the transcribed regions are also fragile for DSBs (Chiarle et al. 2011). In this context, translocation-associated DSBs were enriched at the transcription start sites (TSSs) of active genes but not the inactive genes (Supplemental Fig. S4C). In the PEM-seq libraries from Ku80- or Lig4-deficient CH12F3 cells, translocation levels at TSSs were significantly higher than those from WT cells or Parp1-deficient cells, indicating elevated levels of DSBs in transcribed regions (Fig. 4E).

## Discussion

The off-target activity has been considered to be the main obstacle to clinical applications of CRISPR-Cas9 and similar Cas-associated genome editing toolboxes (Cho et al. 2014; Frock et al. 2015; Tsai et al. 2015; Kim et al. 2019). Recently, more and more abnormal chromosomal structures including large deletions and translocations induced by CRISPR-Cas9 have been observed by different laboratories (Frock et al. 2015; Zuo et al. 2017; Adikusuma et al. 2018; Kosicki et al. 2018; Cullot et al. 2019; Yin et al. 2019; Zuccaro et al. 2020). These inevitable deleterious repair byproducts are generated by endogenous DNA repair pathways and cannot be easily overcome by developed high-fidelity Cas9 variants (Fig. 1D). It has become another dimension of threat to genome stability besides off-targets during genome editing. In order to comprehensively assess DNA repair outcomes during genome editing, we here propose a new *in silico* analysis pipeline PEM-Q with a linear amplification-based sequencing method PEM-seq. Compared to previous CRISPR evaluation assays, PEM-Q is equally well for detecting insertions and deletions but more sensitive for identifying translocations and off-targets (Fig. 1).

Small deletions facilitate the disruption of target genes and are the preferred products of CRISPR-Cas9. However, large deletions may disturb neighbor genes within even hundreds of kb from the target sites (Hu et al. 2014; Frock et al. 2015; Kosicki et al. 2018; Cullot et al. 2019; Yin et al. 2019). We identified thousands of large deletions in each PEM-seq library from mouse or human cells, lymphocytes or embryonic stem cells and found that microhomologies are prevalent at large deletions. Different from deletions, small insertions are predictable as revealed in this study and also described previously (Shou et al. 2018; Chakrabarti et al. 2019). The source of large insertions is mainly some DNA fragments that co-exist in the cell during CRISPR-Cas9 editing, including damaged plasmids or virus backbones (Hanlon et al. 2019; Nguyen Tran et al. 2020; Norris et al. 2020). To suppress deleterious large insertions in clinical applications, DNA-based transfection methods for Cas9 delivery should be avoided.

Translocations occur in one out of hundreds of CRISPR-Cas9-edited cells extrapolated from our findings. Translocations required two simultaneous DSBs that can interact with each other before being fused. In this context, in parallel multiple-gene targeting would induce tremendous translocations between any two target sites. Moreover, recurrent DSBs induced by off-target activity or other physiologic or pathological situations also pose a great threat to genome integrity during genome editing. For instance, translocations induced during V(D)J recombination and class switch recombination usually cause lymphoid tumorigenesis (Alt et al. 2013; Hu et al. 2014; Hu et al. 2015; Zhao et al. 2020). We also showed in this study that the deficiency for NHEJ factor Ku80 or Lig4 leads to a significant increase of large deletions, large insertions, and DSBs around TSSs (Fig. 2H, 3D, and 4E). Therefore, previously developed methods to employ NHEJ inhibitors for promoting HR are not applicable and may pose a great threat to genome integrity during genome editing. Furthermore, besides providing the guidelines for further improving the fidelity of genome editing, PEM-seq also shows great potential to distinguish various DNA repair products in studying DNA repair pathways.

## Methods

### Plasmid construction

All gRNAs used for HEK293T cells and mESCs targeting have been cloned into the double BbsI sites of pX330-vector (Addgene ID 42230). The plasmid used for CH12F3 cell targeting was an optimized vector in which we removed the AAV2 ITR sequence and introduced a mCherry gene with CMV promoter by Gibson assembly into pX330 vector.

### Cell culture and plasmid transfection

The mESCs were cultured in ES-DMEM medium (Millipore) with 15% fetal bovine serum (FBS, ExCell Bio), Penicillin/Streptomycin (Corning), Nucleotides (Millipore), L-Glutamine (Corning), Nonessential Amino Acids (Corning), PD0325901 (Selleck), CHIR99021 (Selleck) and LIF (Millipore) at 37°C with 5% CO_2_. mESCs in 6-cm dishes were transfected with 7.2 μg pX330-Cas9 plus 1.8 μg GFP expression vector by 4D-nucleofector X (Lonza, solution Cytomix, program GC104), then harvested for genomic DNA 3 days after transfection.

The wild-type, Ku80^-/-^, Lig4^-/^, Parp1^-/-^, and AID^-/-^ CH12F3 cells were cultured in RPIM1640 medium (Corning) with 15% Fetal Bovine Serum (FBS, ExCell Bio), HEPES (Corning), Penicillin-Streptomycin (Corning), L-Glutamine (Corning), Nonessential Amino Acids (Corning), Sodium Pyruvate (Corning) and β-Mercaptoethanol (Sigma-Aldrich) at 37°C with 5% CO_2_. Growing CH12F3 cells were transfected with 1.5 μg pX330-Cas9 or pX330-Cas9-mCherry expression vector per million by 4D-nucleofector X (Lonza, solution M1, procedure DN100) and seeding at 0.5 million cells/mL in fresh medium with 1μg/mL anti-CD40, 5 ng/mL IL-4, and 0.5 ng/mL TGF-β. After 72 hrs stimulation, the cells were harvested and genomic DNA was extracted for PEM-seq library construction.

### PEM-seq and 3C-HTGTS

The primers and gRNAs used for library construction are listed in table S2 and table S3, respectively. The PEM-seq libraries were constructed according to the standard procedure described previously (Yin et al. 2019). About 20 μg genome DNA from edited cells were used for each library. Primer control libraries were done with Cas9-infected cells with no gRNA.

The 3C-HTGTS libraries were constructed following the previously described procedures (Jain et al. 2018). Briefly for preparing the 3C-HTGTS libraries, 5-6 million cells were incubated with 1% formaldehyde for 10 min at room temperature and glycine was added to a final concentration of 125 mM to stop the cross-linking reaction. Then cell lysis buffer containing 10mM Tris-HCl (pH 8.0), 10mM NaCl, 0.2% NP-40, 10mM EDTA was used to lysis cell and prepare nuclei. Then the nuclei restriction enzyme (RE) digestion was performed by incubating with 700 units of Dpn II restriction enzyme overnight at 37°C, and the digestion efficiency was checked by DNA gel electrophoresis. Re-ligate the DNA sequence at 16°C for 4 hrs to overnight under dilute conditions. De-crosslink the nuclei by incubating the DNA with Proteinase K at 56°C by rotating overnight. Finally, the purified DNA after RNase A treatment was the “3C templates” and then subsequently prepared the 3C library as the same as PEM-seq library construction.

All the libraries were sequenced by Hiseq.

### PEM-Q analysis

Before PEM-Q analysis, raw reads were pre-processed as we did in the previous method (Yin et al. 2019). We used *cutadapt* (http://cutadapt.readthedocs.io/en/stable/) to remove the universal adapters. Reads ending with QC < 30 were trimmed; remaining reads larger than 25 bp were kept for library demultiplex by *fastq-multx* (https://github.com/brwnj/fastq-multx). Reads after demultiplex were analyzed by PEM-Q in 5 steps.

1. ***reads alignment*** 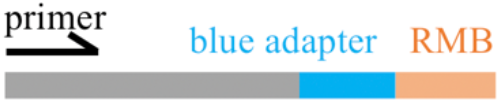 To begin with, R1 and R2 of pair-end reads generated by Hiseq were stitched using *flash 1.2.11* (https://ccb.jhu.edu/software/FLASH/) with default parameters. Then the stitched reads, along with unstitched R1 reads were aligned to reference genome (hg38 for human, mm10 for mouse) by *bwa-mem.* Reads were kept if their alignment start sites were around primer start with an error less than 4 bp. Meanwhile, R2 reads were aligned to the blue adapter, which was used to find random molecular barcode (RMB, equal to unique molecular index) in step 2. Mapped reads with the wrong primer location were discarded in this step.
2. ***RMB extract*** We kept reads with the correct blue adapter allowing at most 2 bp truncation. Then, RMB within 2-bp loss in length were extracted according to blue adapter location. RMB was recorded in a separated file with sequence name (Qname). Reads with multiple tandem adapters were filtered in this step.
3. ***find chimeric alignment*** 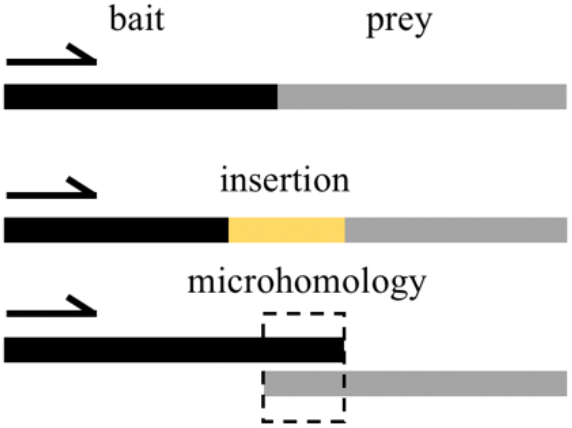 Chimeric reads were reported in SA tag in *bwa-mem.* Sequence aligned to primer was bait while the other side was prey. We then kept reads that only reported one chimeric junction and recorded their information as prey in a tab file. Reads with bait alignment not exceeding 10 bp after primer binding site were discarded. Extra bases between bait and prey were extracted and recorded as insertions. For those without insertions, we identified overlapped bases as microhomology between the end of bait and the start of prey. Reads that did not have chimeric alignment were further analyzed in step 4.
4. ***find indels*** 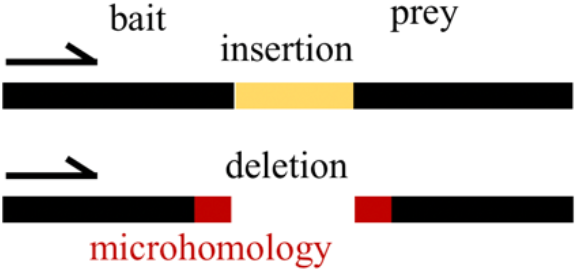 Reads without chimeric alignment were linear alignment. Linear alignment length not exceeding 10 bp after cut-site of CRISPR was discarded. The remaining were processed to find indels. Insertions and deletions were reported by “I” and “D” from CIGAR reported by *bwa-mem.* The same bases at the ends of deletions were identified as microhomology. Substitutions were also aware of PEM-Q and we identified substitutions according to MD tags reported by *bwa-mem*. The remaining reads without chimeric alignment or indels were recorded as germline.
5. ***Classify and deduplicate*** Reads that have both bait and prey aligning to target chromosomes with inserted sequences were classified as insertions. Those without inserted sequences but with a distance between bait and prey no more than 500 kb were classified as deletions in this study. Reads with a distance between bait and prey exceeded 500 kb were classified as intra-chromosomal translocations, while those with prey from other chromosomes were classified as inter-chromosomal translocations. RMB extract in step 2 was relocated to reads according to their sequence name. Within each type of variants we classified, duplicates were removed according to prey’s alignment information including chromosome, strand, junction, and bait end together with RMB.

#### Additional program: Vector (plasmid) analysis

There are two main types of vector integrations as described in the text. One is short vector insertions that the entire inserted fragments can be aligned to the vector backbone. The others with too long inserted fragments are discarded in PEM-Q. However, the second type still has potential large vector integrations. Therefore, we remapped these discarded reads to the genome and then the vector backbone to find missed vector integrations. We used *bwa-mem* to do the alignment with a default seed length of 20 bp.

### Off-target and TSS analysis

Off-target identification was described previously (Yin et al. 2019), using MACS2 *callpeak* and a commonly used criteria. For TSS analysis, we used computeMatrix *(deeptools 3.1.3)* to calculate the signals in RNA Pol II ChIP-seq data, using parameters “-a 50000 -b 50000 -bs 1000”. As for PEM-seq data, we used the same algorithm described before (Zhang et al. 2012) to assign junctions to the nearest TSSs.

## Data access

Sequencing data were deposited into NODE (National Omics Data Encyclopedia, OEP001736) and the PEM-Q pipeline is available at the GitHub site: https://github.com/liumz93/PEM-Q. Other data needed to evaluate the conclusions in the paper are present in the paper and/or the Supplementary Materials. Additional data related to this paper may be requested from the authors.

## Acknowledgement

We thank Dr. Kefei Yu and Dr. Xiong Ji for their generous gifts of Lig4-deficient CH12F3 cells and mESCs. We thank the other lab members for their helpful comments. We thank the Core Facility at the National Center for Protein Sciences, Peking University, particularly the Flow Cytometry Core, for technical help. This work was supported by the National Key R&D Program of China (2017YFA0506700 to J.H. and F.-L. M.) and the NSFC grant (31771485 to J.H.).

## Author contributions

M.L., W.Z., C.X., and J.H. conceived and designed the study; M.L. developed the PEM-Q pipeline and J.L. was involved in verifying the pipeline; W.Z., C.X., J.Y., and Y.S. performed the experiments; Y.S. and F.-L. M. generated CH12F3 cell mutants; M.L., W.Z., C.X. C.A., F.-L. M., and J.H. participated in analyzing the sequencing data; M.L., W.Z., C.X. and J.H. wrote the paper.

## References

Adikusuma F, Piltz S, Corbett MA, Turvey M, McColl SR, Helbig KJ, Beard MR, Hughes J, Pomerantz RT, Thomas PQ. 2018. Large deletions induced by Cas9 cleavage. Nature 560:E8–E9.

Alt FW, Zhang Y, Meng FL, Guo C, Schwer B. 2013. Mechanisms of programmed DNA lesions and genomic instability in the immune system. Cell 152:417–429.

Anzalone AV, Koblan LW, Liu DR. 2020. Genome editing with CRISPR-Cas nucleases, base editors, transposases and prime editors. Nat Biotechnol 38:824–844.

Chakrabarti AM, Henser-Brownhill T, Monserrat J, Poetsch AR, Luscombe NM, Scaffidi P. 2019. Target-Specific Precision of CRISPR-Mediated Genome Editing. Mol Cell 73:699–713 e696.

Chang HHY, Pannunzio NR, Adachi N, Lieber MR. 2017. Non-homologous DNA end joining and alternative pathways to double-strand break repair. Nat Rev Mol Cell Biol 18:495–506.

Chiarle R, Zhang Y, Frock RL, Lewis SM, Molinie B, Ho YJ, Myers DR, Choi VW, Compagno M, Malkin DJ et al. 2011. Genome-wide translocation sequencing reveals mechanisms of chromosome breaks and rearrangements in B cells. Cell 147:107–119.

Cho SW, Kim S, Kim Y, Kweon J, Kim HS, Bae S, Kim JS. 2014. Analysis of off-target effects of CRISPR/Cas-derived RNA-guided endonucleases and nickases. Genome Res 24:132–141.

Chu VT, Weber T, Wefers B, Wurst W, Sander S, Rajewsky K, Kuhn R. 2015. Increasing the efficiency of homology-directed repair for CRISPR-Cas9-induced precise gene editing in mammalian cells. Nat Biotechnol 33:543–548.

Cong L, Ran FA, Cox D, Lin S, Barretto R, Habib N, Hsu PD, Wu X, Jiang W, Marraffini LA et al. 2013. Multiplex genome engineering using CRISPR/Cas systems. Science 339:819–823.

Cullot G, Boutin J, Toutain J, Prat F, Pennamen P, Rooryck C, Teichmann M, Rousseau E, Lamrissi-Garcia I, Guyonnet-Duperat V et al. 2019. CRISPR-Cas9 genome editing induces megabase-scale chromosomal truncations. Nat Commun 10: 1136.

Doudna JA. 2020. The promise and challenge of therapeutic genome editing. Nature 578:229–236.

Frock RL, Hu J, Meyers RM, Ho YJ, Kii E, Alt FW. 2015. Genome-wide detection of DNA doublestranded breaks induced by engineered nucleases. Nat Biotechnol 33:179–186.

Hanlon KS, Kleinstiver BP, Garcia SP, Zaborowski MP, Volak A, Spirig SE, Muller A, Sousa AA, Tsai SQ, Bengtsson NE et al. 2019. High levels of AAV vector integration into CRISPR-induced DNA breaks. Nat Commun 10: 4439.

Hendriks D, Clevers H, Artegiani B. 2020. CRISPR-Cas Tools and Their Application in Genetic Engineering of Human Stem Cells and Organoids. Cell Stem Cell 27:705–731.

Hu J, Meyers RM, Dong J, Panchakshari RA, Alt FW, Frock RL. 2016. Detecting DNA double-stranded breaks in mammalian genomes by linear amplification-mediated high-throughput genome-wide translocation sequencing. Nat Protoc 11:853–871.

Hu J, Tepsuporn S, Meyers RM, Gostissa M, Alt FW. 2014. Developmental propagation of V(D)J recombination-associated DNA breaks and translocations in mature B cells via dicentric chromosomes. Proc Natl Acad Sci U S A 111:10269–10274.

Hu J, Zhang Y, Zhao L, Frock RL, Du Z, Meyers RM, Meng FL, Schatz DG, Alt FW. 2015. Chromosomal Loop Domains Direct the Recombination of Antigen Receptor Genes. Cell 163:947–959.

Hu Z, Shi Z, Guo X, Jiang B, Wang G, Luo D, Chen Y, Zhu YS. 2018. Ligase IV inhibitor SCR7 enhances gene editing directed by CRISPR-Cas9 and ssODN in human cancer cells. Cell Biosci 8: 12.

Jain S, Ba Z, Zhang Y, Dai HQ, Alt FW. 2018. CTCF-Binding Elements Mediate Accessibility of RAG Substrates During Chromatin Scanning. Cell 174:102–116 e114.

Jinek M, Chylinski K, Fonfara I, Hauer M, Doudna JA, Charpentier E. 2012. A programmable dual-RNA-guided DNA endonuclease in adaptive bacterial immunity. Science 337:816–821.

Jinek M, East A, Cheng A, Lin S, Ma E, Doudna J. 2013. RNA-programmed genome editing in human cells. Elife 2: e00471.

Kim D, Kim DE, Lee G, Cho SI, Kim JS. 2019. Genome-wide target specificity of CRISPR RNA-guided adenine base editors. Nat Biotechnol 37:430–435.

Kosicki M, Tomberg K, Bradley A. 2018. Repair of double-strand breaks induced by CRISPR-Cas9 leads to large deletions and complex rearrangements. Nat Biotechnol 36:765–771.

Ling X, Xie B, Gao X, Chang L, Zheng W, Chen H, Huang Y, Tan L, Li M, Liu T. 2020. Improving the efficiency of precise genome editing with site-specific Cas9-oligonucleotide conjugates. Sci Adv 6: eaaz0051.

Liu X, Liu T, Shang Y, Dai P, Zhang W, Lee BJ, Huang M, Yang D, Wu Q, Liu LD et al. 2020. ERCC6L2 promotes DNA orientation-specific recombination in mammalian cells. Cell Res 30:732–744.

Mali P, Yang L, Esvelt KM, Aach J, Guell M, DiCarlo JE, Norville JE, Church GM. 2013. RNA-guided human genome engineering via Cas9. Science 339:823–826.

Maruyama T, Dougan SK, Truttmann MC, Bilate AM, Ingram JR, Ploegh HL. 2015. Increasing the efficiency of precise genome editing with CRISPR-Cas9 by inhibition of nonhomologous end joining. Nat Biotechnol 33:538–542.

Nguyen GN, Everett JK, Kafle S, Roche AM, Raymond HE, Leiby J, Wood C, Assenmacher CA, Merricks EP, Long CT et al. 2021. A long-term study of AAV gene therapy in dogs with hemophilia A identifies clonal expansions of transduced liver cells. Nat Biotechnol 39:47–55.

Nguyen Tran MT, Mohd Khalid MKN, Wang Q, Walker JKR, Lidgerwood GE, Dilworth KL, Lisowski L, Pebay A, Hewitt AW. 2020. Engineering domain-inlaid SaCas9 adenine base editors with reduced RNA off-targets and increased on-target DNA editing. Nat Commun 11: 4871.

Norris AL, Lee SS, Greenlees KJ, Tadesse DA, Miller MF, Lombardi HA. 2020. Template plasmid integration in germline genome-edited cattle. Nat Biotechnol 38:163–164.

Owens DDG, Caulder A, Frontera V, Harman JR, Allan AJ, Bucakci A, Greder L, Codner GF, Hublitz P, McHugh PJ et al. 2019. Microhomologies are prevalent at Cas9-induced larger deletions. Nucleic Acids Res 47:7402–7417.

Pawluk A, Davidson AR, Maxwell KL. 2018. Anti-CRISPR: discovery, mechanism and function. Nat Rev Microbiol 16:12–17.

Pinello L, Canver MC, Hoban MD, Orkin SH, Kohn DB, Bauer DE, Yuan GC. 2016. Analyzing CRISPR genome-editing experiments with CRISPResso. Nat Biotechnol 34:695–697.

Rees HA, Yeh WH, Liu DR. 2019. Development of hRad51-Cas9 nickase fusions that mediate HDR without double-stranded breaks. Nat Commun 10: 2212.

Riesenberg S, Maricic T. 2018. Targeting repair pathways with small molecules increases precise genome editing in pluripotent stem cells. Nat Commun 9: 2164.

Sfeir A, Symington LS. 2015. Microhomology-Mediated End Joining: A Back-up Survival Mechanism or Dedicated Pathway? Trends Biochem Sci 40:701–714.

Shou J, Li J, Liu Y, Wu Q. 2018. Precise and Predictable CRISPR Chromosomal Rearrangements Reveal Principles of Cas9-Mediated Nucleotide Insertion. Mol Cell 71:498–509 e494.

Truong LN, Li Y, Shi LZ, Hwang PY, He J, Wang H, Razavian N, Berns MW, Wu X. 2013. Microhomology-mediated End Joining and Homologous Recombination share the initial end resection step to repair DNA double-strand breaks in mammalian cells. Proc Natl Acad Sci U S A 110:7720–7725.

Tsai SQ, Zheng Z, Nguyen NT, Liebers M, Topkar VV, Thapar V, Wyvekens N, Khayter C, Iafrate AJ, Le LP et al. 2015. GUIDE-seq enables genome-wide profiling of off-target cleavage by CRISPR-Cas nucleases. Nat Biotechnol 33:187–197.

Wang D, Zhang F, Gao G. 2020. CRISPR-Based Therapeutic Genome Editing: Strategies and In Vivo Delivery by AAV Vectors. Cell 181:136–150.

Yeh CD, Richardson CD, Corn JE. 2019. Advances in genome editing through control of DNA repair pathways. Nat Cell Biol 21:1468–1478.

Yin J, Liu M, Liu Y, Wu J, Gan T, Zhang W, Li Y, Zhou Y, Hu J. 2019. Optimizing genome editing strategy by primer-extension-mediated sequencing. Cell Discov 5: 18.

Zhang Y, McCord RP, Ho YJ, Lajoie BR, Hildebrand DG, Simon AC, Becker MS, Alt FW, Dekker J. 2012. Spatial organization of the mouse genome and its role in recurrent chromosomal translocations. Cell 148:908–921.

Zhao B, Rothenberg E, Ramsden DA, Lieber MR. 2020. The molecular basis and disease relevance of non-homologous DNA end joining. Nat Rev Mol Cell Biol 21:765–781.

Zuccaro MV, Xu J, Mitchell C, Marin D, Zimmerman R, Rana B, Weinstein E, King RT, Palmerola KL, Smith ME et al. 2020. Allele-Specific Chromosome Removal after Cas9 Cleavage in Human Embryos. Cell 183:1650–1664 e1615.

Zuo E, Huo X, Yao X, Hu X, Sun Y, Yin J, He B, Wang X, Shi L, Ping J et al. 2017. CRISPR/Cas9-mediated targeted chromosome elimination. Genome Biol 18: 224.

